# Maternal *Med12* safeguards trophoblast pluripotency and placental development

**DOI:** 10.1101/2024.11.01.621455

**Authors:** Michelle M. Halstead, Jyoti Goad, Amina Khan, Rutuja Deshmukh, Aleksandar Rajkovic

## Abstract

For a brief but critical period post-fertilization, the mammalian embryo is entirely dependent on maternal products inherited from the oocyte. Previous research showed that oocyte-specific loss of *Med12*, an X-linked gene and Mediator complex subunit, leads to female sterility despite normal folliculogenesis and ovulation. Here, we show that loss of maternal *Med12* has minimal effect on the oocyte transcriptome and does not manifest in embryonic lethality until post-implantation. Implants derived from *Med12*-null oocytes demonstrate abnormal placentation at E9.5, with an overabundance of trophoblast giant cells (TGC). This phenotype associates with downregulation of trophoblast pluripotency markers (e.g. *Cdx2*) and activation of drivers of TGC identity (e.g. *Stra13*) in the E7.5 extraembryonic ectoderm, revealing a previously undescribed role for *Med12* in trophoblast pluripotency maintenance. Notably, we find consistently low *Med12* expression in embryos derived from *Med12*-null oocytes, potentially due to programmed paternal X chromosome inactivation (XCI). To isolate the consequences of maternal *Med12* depletion, we introduced an autosomal *Med12* transgene and show that embryonic expression of the transgene rescues development of *Med12*-null oocytes. We conclude that oocyte-specific deletion of *Med12* produces a maternal-zygotic double knock-out in extraembryonic tissues due to paternal XCI, leading to loss of pluripotency in the trophoblast, placental malformation, and embryonic death.

## Introduction

Early pregnancy loss (EPL) remains a significant challenge in human reproductive medicine, affecting about 10% of clinically recognized pregnancies (Zinaman et al., 1996). An estimated 80% of miscarriages occur in the first trimester of pregnancy (X. Wang et al., 2003), often before the person knows they are pregnant. However, very little is known about the mechanisms that govern successful development during early embryogenesis. Maternal effect genes (MEG), which encode oocyte-derived factors that are dispensable for oogenesis but essential for embryogenesis, are of particular interest, as EPL driven by MEG dysregulation is determined solely by the mother’s genotype. MEG guide embryonic development before the onset of embryonic transcription, regulating crucial processes such as zygotic genome activation (ZGA), epigenetic reprogramming, and cell differentiation (Mitchell, 2022). Disruption of MEG can profoundly impact embryonic development, potentially leading to EPL or birth defects. However, relatively few MEGs have been described in mammals.

Using Zp3Cre-mediated recombination, we previously demonstrated that oocyte-specific depletion of Mediator complex subunit 12 (*Med12*) leads to female sterility without impacting folliculogenesis or ovulation (X. Wang et al., 2017). This finding suggests that *Med12* may play a novel role as an MEG in mice. The Mediator complex is a critical determinant of overall genome organization and gene expression, acting as a bridge between gene-specific transcription factors bound to enhancer elements and the RNA polymerase II transcription machinery at gene promoters (Kagey et al., 2010; Phillips-Cremins et al., 2013). Structurally, the Mediator complex is organized into three core modules—head, middle, and tail—comprising 26 subunits, along with a detachable kinase module consisting of Med12, Med13, Cyclin C, and CDK8, which regulates the function of the core Mediator (Harper & Taatjes, 2018; Knuesel et al., 2009; Nozawa et al., 2017; Tsai et al., 2017). Med12 is essential for the activation of CDK8 within the kinase module, which can modulate the interaction of Mediator with RNA polymerase II, thereby influencing transcription (Galli et al., 2015; Jaeger et al., 2020; Soutourina, 2018; Takahashi et al., 2011; G. Wang et al., 2005) and chromatin architecture (Aranda-Orgilles et al., 2016; El Khattabi et al., 2019; Haarhuis et al., 2022; Kagey et al., 2010; Phillips-Cremins et al., 2013; Whyte et al., 2013) in a context-dependent manner. Recent evidence also indicates that Med12-dependent CDK8 activation plays a key role in various signaling pathways, including those related to nuclear receptors, Wnt, and Sonic Hedgehog, and that the Med12-containing CDK8 subcomplex can function independently of the core Mediator complex (Aranda-Orgilles et al., 2016).

Previous studies established that *Med12* is required for embryonic development in mice, with its loss compromising Wnt signaling and gastrulation in embryos generated via tetraploid complementation with *Med12*-null stem cells (Rocha et al., 2010). However, the role of *Med12* in the preimplantation embryo and extraembryonic tissues has not been investigated, and it remains unclear when and how the loss of oocyte-derived *Med12* leads to embryonic arrest and female sterility. To investigate the role of maternal *Med12* in the oocyte and early embryo, we ablated *Med12* from primary follicles via Zp3Cre-mediated recombination and investigated the molecular signatures of the resulting oocytes and embryos. As *Med12* is an X-linked gene, we also investigated the impact of X chromosome inactivation (XCI) on embryonic *Med12* expression. We find that the paternal *Med12* allele is generally inactive in embryos, such that oocyte-specific deletion of *Med12* produces a maternal-zygotic double knock-out in extraembryonic tissues. Consequently, loss of *Med12* leads to placental malformation and downregulation of pluripotency markers in the extraembryonic ectoderm, implicating *Med12* in trophoblast pluripotency maintenance. Finally, embryonic expression of an autosomal *Med12* transgene rescues development of *Med12*-null oocytes, suggesting that the requirement for oocyte-derived *Med12* is rooted in the rodent-specific phenomenon of paternal XCI.

## Results

### Med12 does not regulate transcription in the oocyte

Oocyte-specific knock-out of *Med12* did not affect folliculogenesis or ovulation (X. Wang et al., 2017). However, the molecular signature of *Med12*-ablated oocytes was not assessed. Considering the fundamental role of *Med12* in transcriptional regulation, the lack of *Med12* might alter the oocyte transcriptome and thereby compromise embryonic development, even if ovarian function remains unperturbed.

As reported previously (X. Wang et al., 2017; Wasson et al., 2016), *Med12* was ablated from primary follicles via Zp3Cre-mediated recombination (Figure 1A). As expected, *Med12* transcripts were significantly reduced by about 20-fold in germinal vesicle (GV) and metaphase II (MII) stage oocytes from *Zp3Cre;Med12^fl/fl^*dams compared to *Med12^fl/fl^* control dams (Figure 1B). To determine the consequences of *Med12* depletion on transcription regulation in the oocyte, post-natal day 12 (D12), GV, and MII oocytes were collected from mutant (*Zp3Cre;Med12^fl/fl^*) and control (*Med12^fl/fl^*) dams for bulk RNA-seq (Figure 1C, Supplementary Figure 1). Despite the core role of the Mediator complex in transcription regulation, *Med12* deficiency had very little effect on the oocyte transcriptome. Only 33, 19, and 10 genes were dysregulated (adjusted *p* < 0.01) in D12, GV, and MII stage *Med12*-deficient oocytes, respectively (Figure 1D, Supplementary Data 1). Of these dysregulated genes, *Mt1* was upregulated in both D12 and GV mutant oocytes; however, upregulation of *Mt1* may indirectly result from Zp3Cre transgene expression (Wasson et al., 2016), rather than downregulation of *Med12*. Overall, *Med12* does not appear to play an important role in transcriptional regulation in the oocyte.

**Figure 1.**
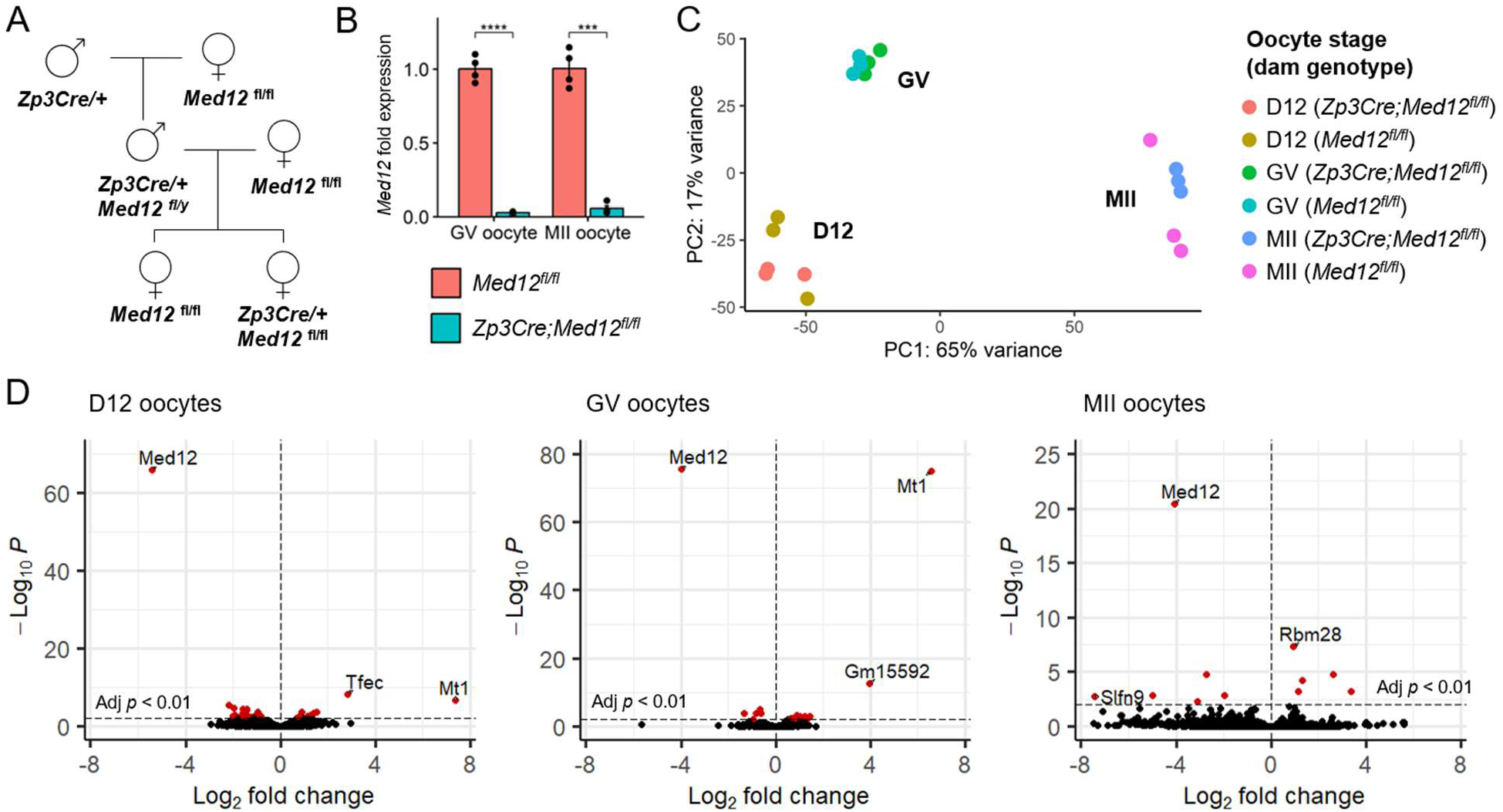
*Med12* ablation does not affect the oocyte transcriptome. (A) Breeding scheme to obtain control (*Med12^fl/fl^*) and mutant (*Zp3Cre;Med12^fl/fl^*) dams. (B) Relative abundance ± S.E.M. of *Med12* in germinal vesicle (GV) and metaphase II (MII) oocytes by RT-qPCR. Calculated by the ΔΔCt method using the reference gene *Gapdh* (n=4 replicates per genotype/stage). One-tailed unpaired Student’s t-test (***p<0.001, ****p<1e-4). (C) Principal components analysis of growing (day 12; D12), GV and MII oocyte transcriptomes from *Med12^fl/fl^* and *Zp3Cre;Med12^fl/fl^* dams. Based on top 5,000 genes with highest variable expression. (D) Differential gene expression in oocytes from control versus mutant dams. Differentially expressed genes colored red (adjusted *p* < 0.01).

**Supplementary Figure 1.**
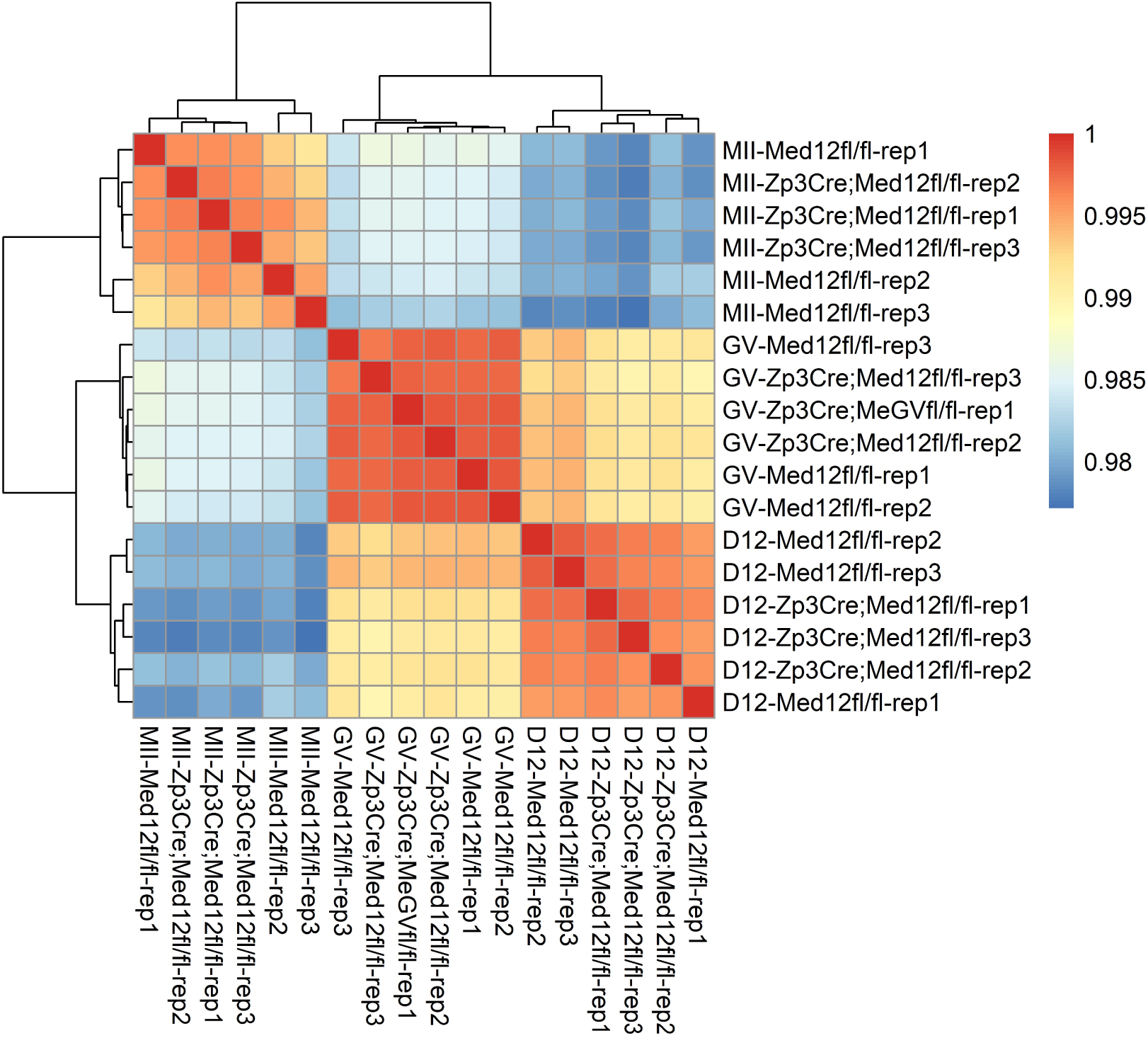
Pearson correlation of oocyte transcriptomes based on bulk RNA-seq. Samples grouped by unsupervised hierarchical clustering. Labels indicate oocyte stage (D12, GV, or MII), dam genotype (*Med12^fl/fl^* or *Zp3Cre;Med12^fl/fl^*) and replicate.

### Maternal Med12 is required for trophoblast development

Transcription was not dysregulated in *Med12*-deficient oocytes, suggesting that maternal *Med12* is not required until after fertilization. To understand when and how loss of maternal *Med12* leads to female sterility, control (*Med12^fl/fl^*) and mutant (*Zp3Cre;Med12^fl/fl^*) dams were timed-mated with wild-type stud males and uteri were collected between E7.5 and E12.5. Similar numbers of implantation sites were observed in control and mutant dams (Supplementary Figure 2). At E9.5, there were no gross morphological differences in implants between control and mutant dams; however, by E12.5 hemorrhagic sites and embryo reabsorption were observed in mutant dams (Figure 2A). As early as E8.5, implants were significantly smaller in mutant dams, and by E12.5, control implants were 10 times larger than mutant implants (Figure 2B). Histological analysis of E7.5 mutant implants did not show any morphological abnormalities, but by E9.5, mutant implants had abnormally high numbers of trophoblast giant cells (TGC) and diminished labyrinth and spongiotrophoblast layers (Figure 2C). This abnormal placental development could explain the diminished embryonic growth and eventual resorption by E12.5.

**Supplementary Figure 2.**
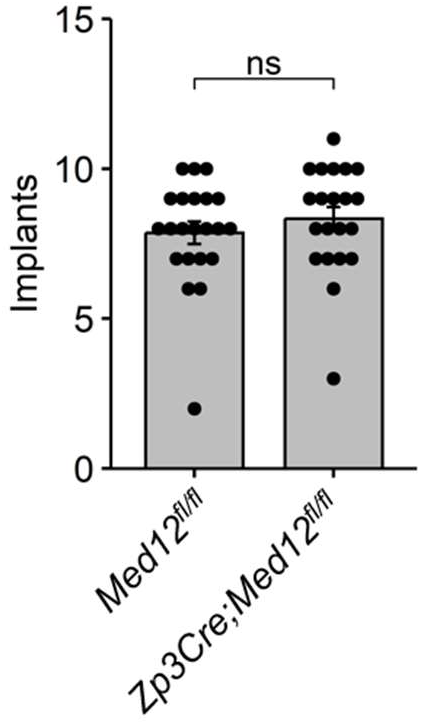
Number of implants observed in control and mutant dams. Uteri collected from control (*Med12^fl/fl^*) and mutant (*Zp3Cre;Med12^fl/fl^*) dams timed mated with wild-type stud males. Uteri collected at E6.5 (n=1 control, n=1 mutant), E7.5 (n=7 control, n=8), E8.5 (n=5 control, n=2 mutant), E9.5 (n=6 control, n=8 mutant) and E12.5 (n=2 control, n=4 mutant). Average implants ± S.E.M. Two-tailed unpaired Student’s t-test p=0.391.

**Figure 2.**
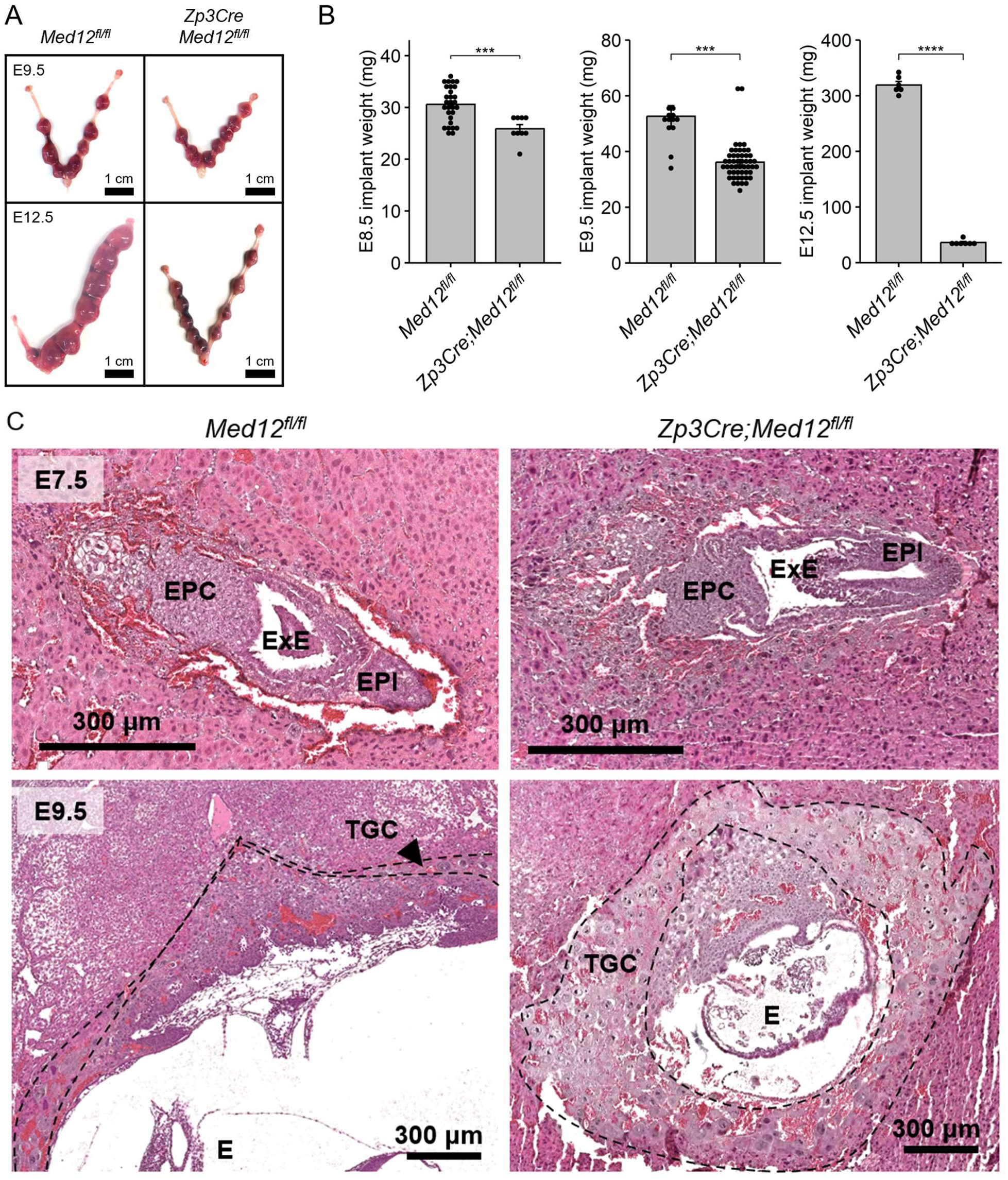
Maternal *Med12* is required for trophoblast development. Uteri collected between E7.5 and E12.5 from control (*Med12^fl/fl^*) and mutant (*Zp3Cre;Med12^fl/fl^*) females mated with wild-type males. (A) Gross analysis of embryo implants at E9.5 and E12.5, showing hemorrhagic sites and embryo reabsorption in mutant dams at E12.5. (B) Weight of individual embryo implants ± S.E.M. at E8.5 (n=29 from control dams; n=9 from mutant dams), E9.5 (n=13 from control dams; n=50 from mutant dams), and E12.5 (n=6 from control dams; n=7 from mutant dams). Two-tailed unpaired Student’s t-test (*** p< 0.001, **** p<1e-4). (C) Histological analysis of placenta and embryo implants from control and mutant dams at E7.5 and E9.5. EPC, ectoplacental cone; ExE, extraembryonic ectoderm; EPI, epiblast; TGC, trophoblast giant cells; E, embryo.

The morphological abnormalities observed in E9.5 mutant embryos likely reflect earlier disruptions to transcription regulation in the lineages that give rise to TGC and other placental tissues. To understand how the loss of maternal *Med12* disrupts placental development, individual E7.5 implants were collected from control and mutant dams for bulk RNA-seq. Each implant was dissected into the epiblast (EPI), extraembryonic ectoderm (ExE), and ectoplacental cone (EPC) (Figure 3A). The sex of each implant was determined by expression of either *Xist* (involved in X chromosome inactivation) or *Ddx3y* (Y-linked gene) (Supplementary Figure 3A). *Med12* was downregulated (adjusted *p* < 0.01) in all tissues of embryos from mutant dams, regardless of embryonic sex (Supplementary Figure 3B). The presence of any *Med12* transcripts in male embryos from mutant dams, which should be null for *Med12* (*Med12Δ/y*), could indicate some contamination from maternal decidua. The consistently low levels of *Med12* in male (*Med12Δ/y*) and female (*Med12Δ/+*) embryos from mutant dams suggests that *Med12* expression from the paternal X chromosome in female embryos is insufficient to restore *Med12* levels to normal, although the expression in epiblast was slightly higher in female than male embryos. Overall, E7.5 samples clearly clustered by embryonic tissue of origin, but did not clearly segregate by sex, nor by maternal genotype (Figure 3B-C). Pairwise comparisons between embryos from mutant dams (lacking maternal *Med12*) and controls were conducted for each embryonic tissue and sex (Table 1, Supplementary Figure 4, Supplementary Data 2). Loss of maternal *Med12* led to dysregulation of core cellular processes, including metabolism, translation, and transcription (Figure 4, Supplementary Figure 5). Consistent with previous reports of *Med12* knock-out embryos generated through tetraploid complementation, depletion of *Med12* in the epiblast disrupted Wnt signaling and somitogenesis (Rocha et al., 2010). Genes involved in cell differentiation, proliferation, migration, and cytoskeletal organization were broadly dysregulated in all E7.5 tissues of *Med12*-depleted embryos (Figure 4). The ExE, which gives rise to specialized placental cell types, was particularly impacted. Markers of trophoblast pluripotency (*Cdx2*, *Esrrb*, *Fgfr2*, *Eomes, Elf5*) were downregulated in the ExE of mutant embryos (Figure 5A), whereas drivers of giant cell identity (*Stra13*, *Ppard, Plet1*) were upregulated in (Figure 5B). We confirmed the downregulation of CDX2 at the protein level in mutant embryos by immunofluorescence (Figure 5C). These data suggest that loss of maternal *Med12* disrupts pluripotency maintenance in the ExE of E7.5 embryos, accelerating differentiation into TGC, which is consistent with the overabundance of TGC at E9.5 (Figure 2C). The infertility of *Zp3Cre;Med12^fl/fl^* mutant dams can therefore be traced to compromised pluripotency maintenance in the peri-implantation trophoblast.

**Figure 3.**
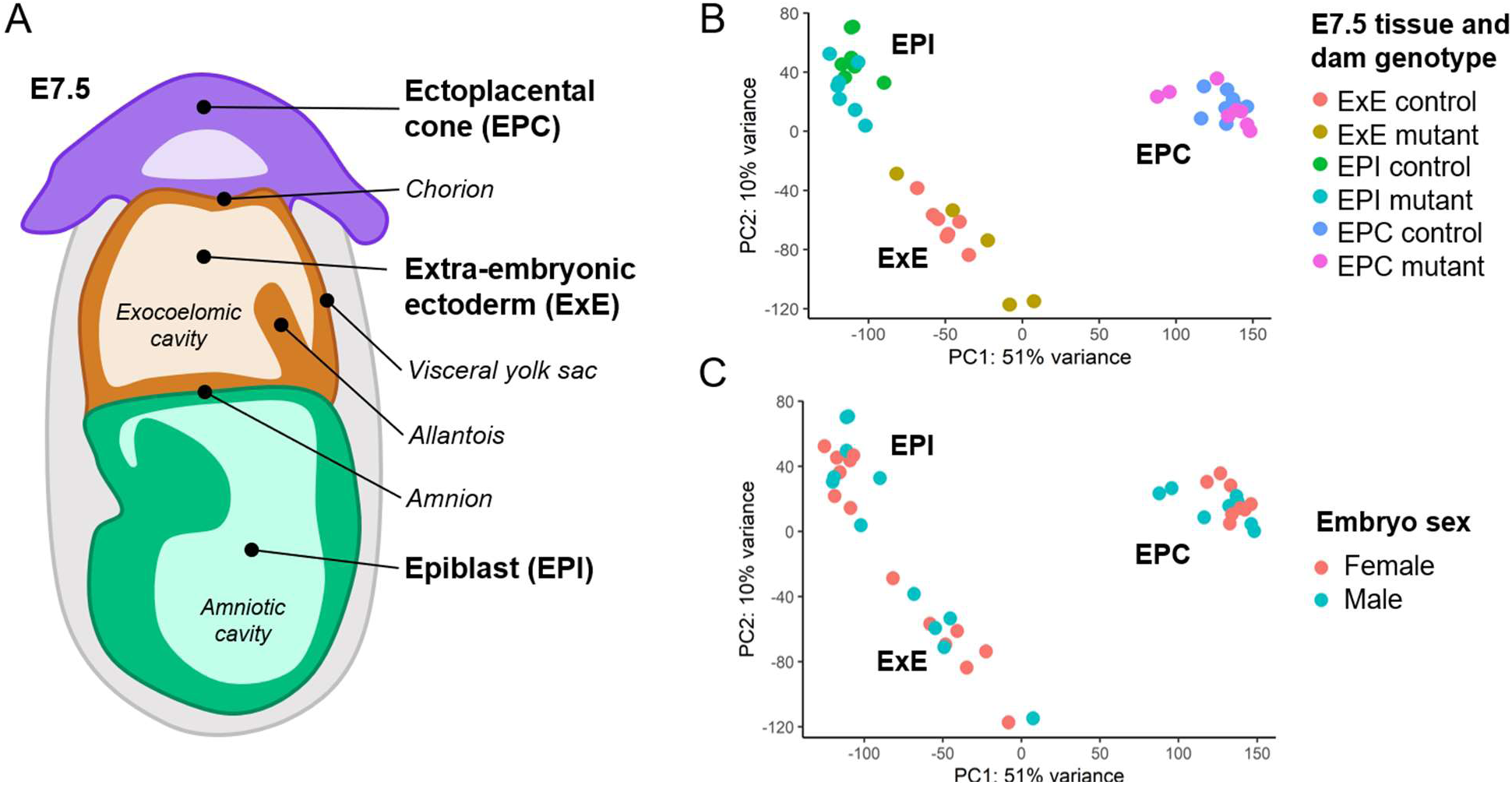
RNA-seq of E7.5 embryos lacking maternal *Med12*. (A) Representation of an E7.5 embryo. (B,C) Principal components analysis of E7.5 transcriptomes, based on top 5,000 genes with highest variable expression. Samples colored based on (B) E7.5 tissue and dam genotype (control, *Med12^fl/fl^*; mutant, *Zp3Cre;Med12^fl/fl^*) or (C) sex of the embryo.

**Supplementary Figure 3.**
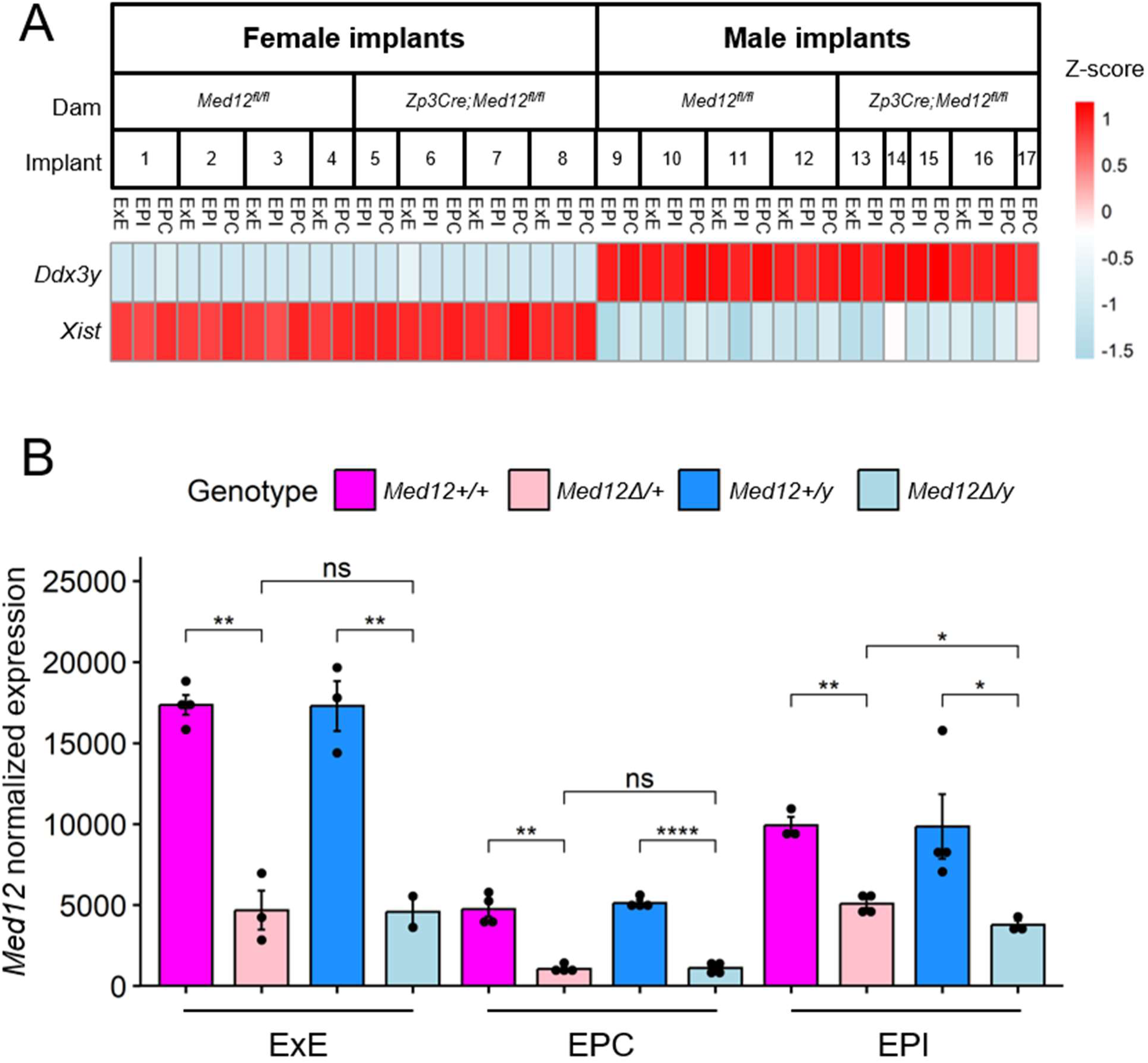
Validation of *Med12* knock down and embryo sex. (A) Heatmap of gene expression Z-scores computed for *Ddx3y* and *Xist*, used to identify E7.5 embryo samples as male or female, respectively. (B) Normalized expression ± S.E.M. of *Med12* in E7.5 samples. Two-tailed unpaired Student’s t-test, adjusted by the Holm-Bonferroni method (ns p≥0.05; * *p*<0.05; ** *p*<0.01; *** *p*<0.001; **** *p*<1e-4).

**Supplementary Figure 4.**
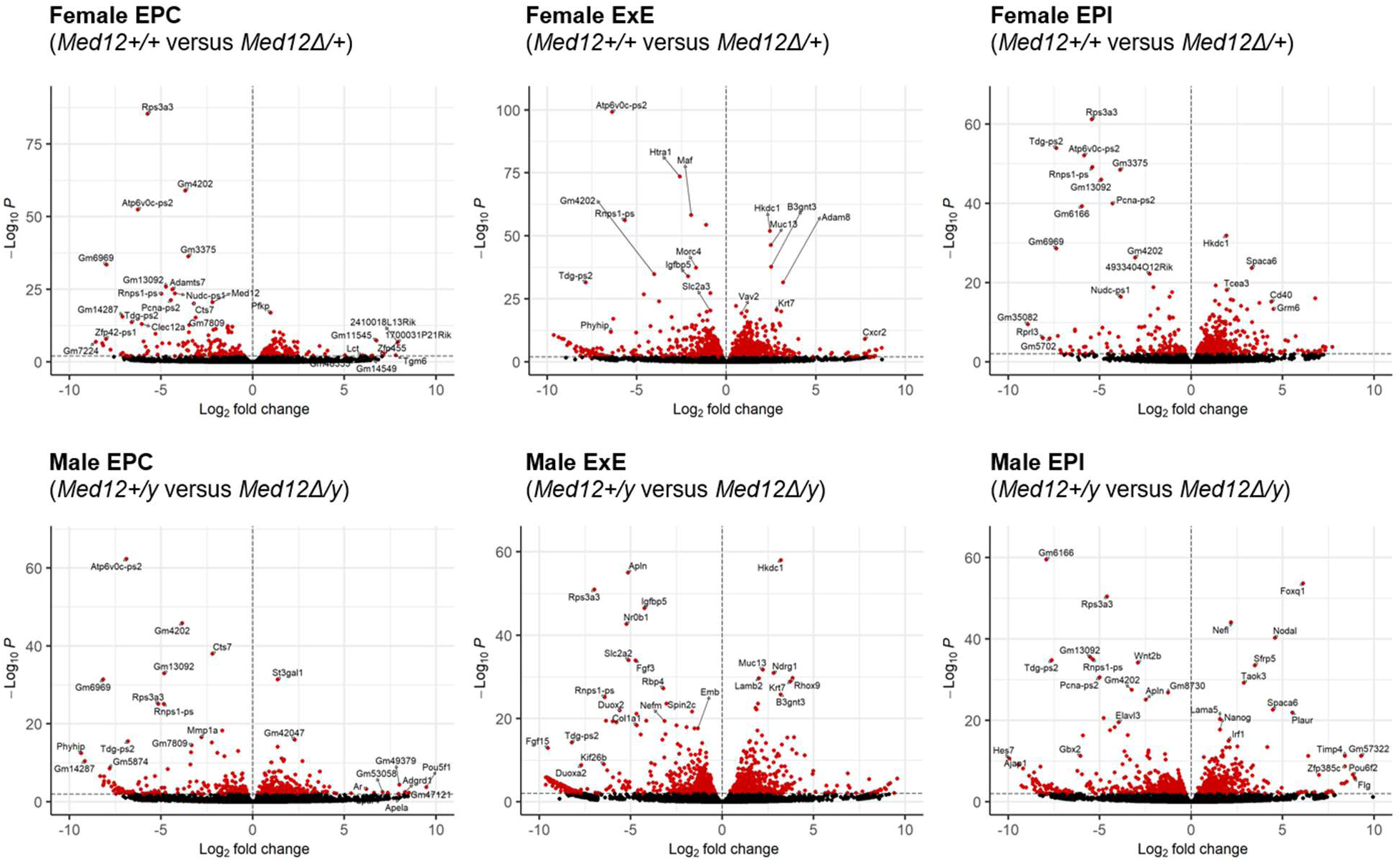
Differentially expressed genes (DEG) between E7.5 embryos from control (*Med12^fl/fl^*) and mutant (*Zp3Cre;Med12^fl/fl^*) dams. Pairwise comparisons conducted for each embryonic tissue (EPC, ExE, and EPI) and sex. DEG colored red (adjusted *p* < 0.01).

**Figure 4.**
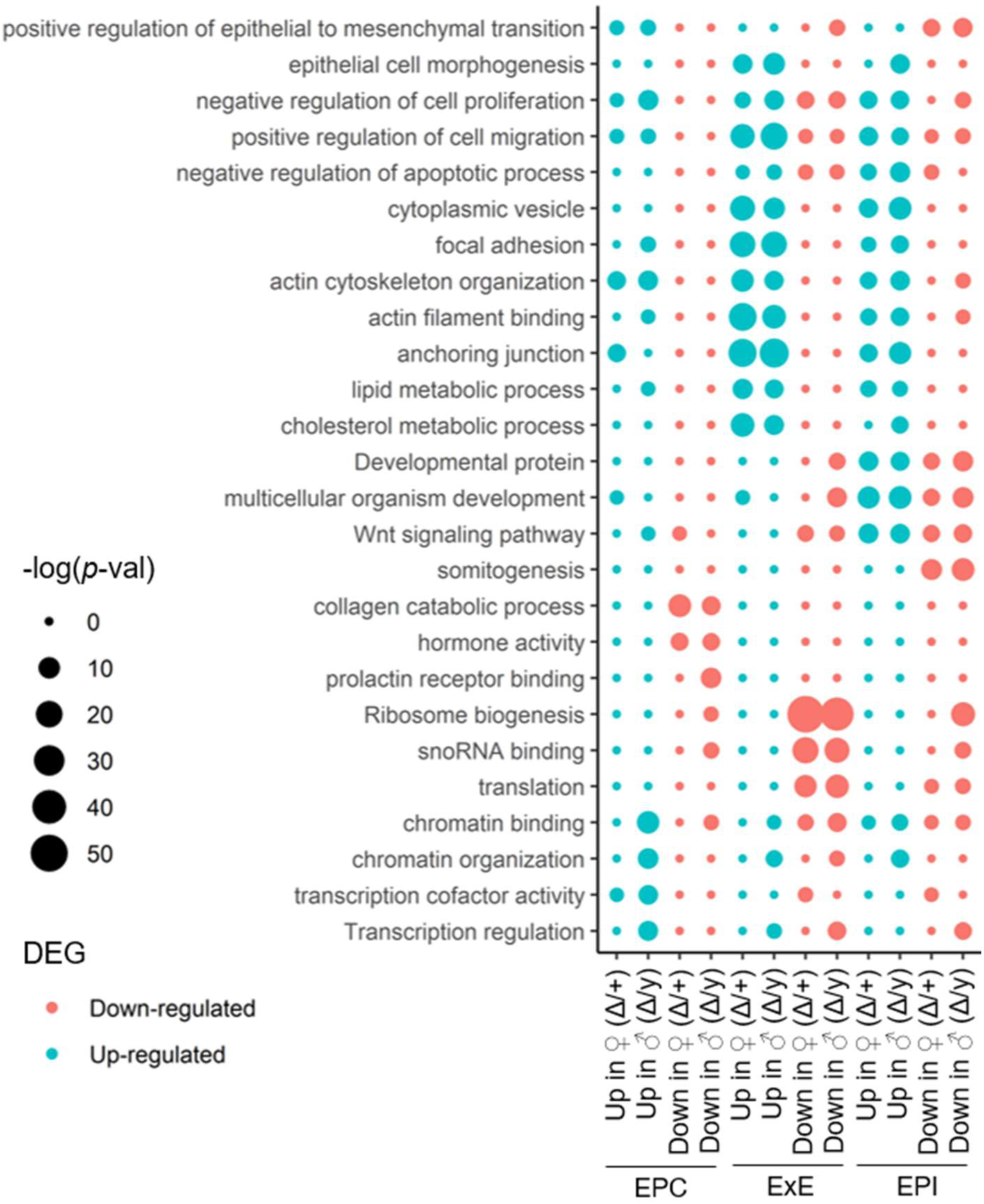
Loss of maternal *Med12* disrupts differentiation, cytoskeletal organization, metabolism, and transcription regulation in E7.5 embryos. Functional enrichment of differentially expressed genes (DEG) that were up-or down-regulated in E7.5 implants collected from mutant (*Zp3Cre;Med12^fl/fl^*) dams. DEG sets were identified separately for each embryo sex and E7.5 tissue. Point size reflects –log(*p*), calculated using the DAVID webtool.

**Supplementary Figure 5.**
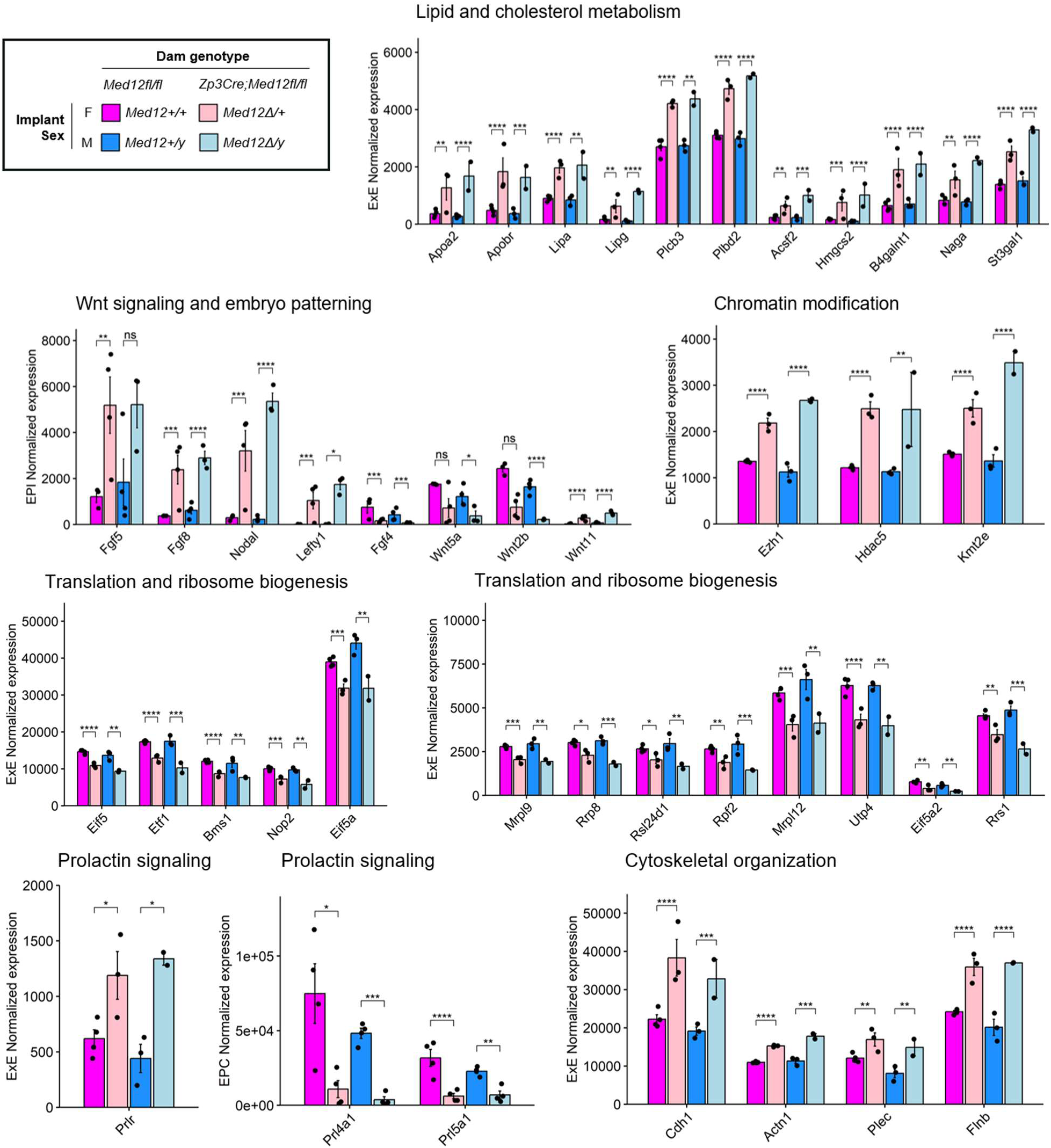
Normalized expression ± S.E.M. of genes involved in lipid and cholesterol metabolism in ExE, Wnt signaling and embryo patterning in EPI, chromatin modification in ExE, translation and ribosome biogenesis in ExE, prolactin signaling in EPC and ExE, and cytoskeletal organization in ExE. Adjusted *p*-values based on differential expression analyses (ns p≥0.05; * *p*<0.05; ** *p*<0.01; *** *p*<0.001; **** *p*<1e-4).

**Figure 5.**
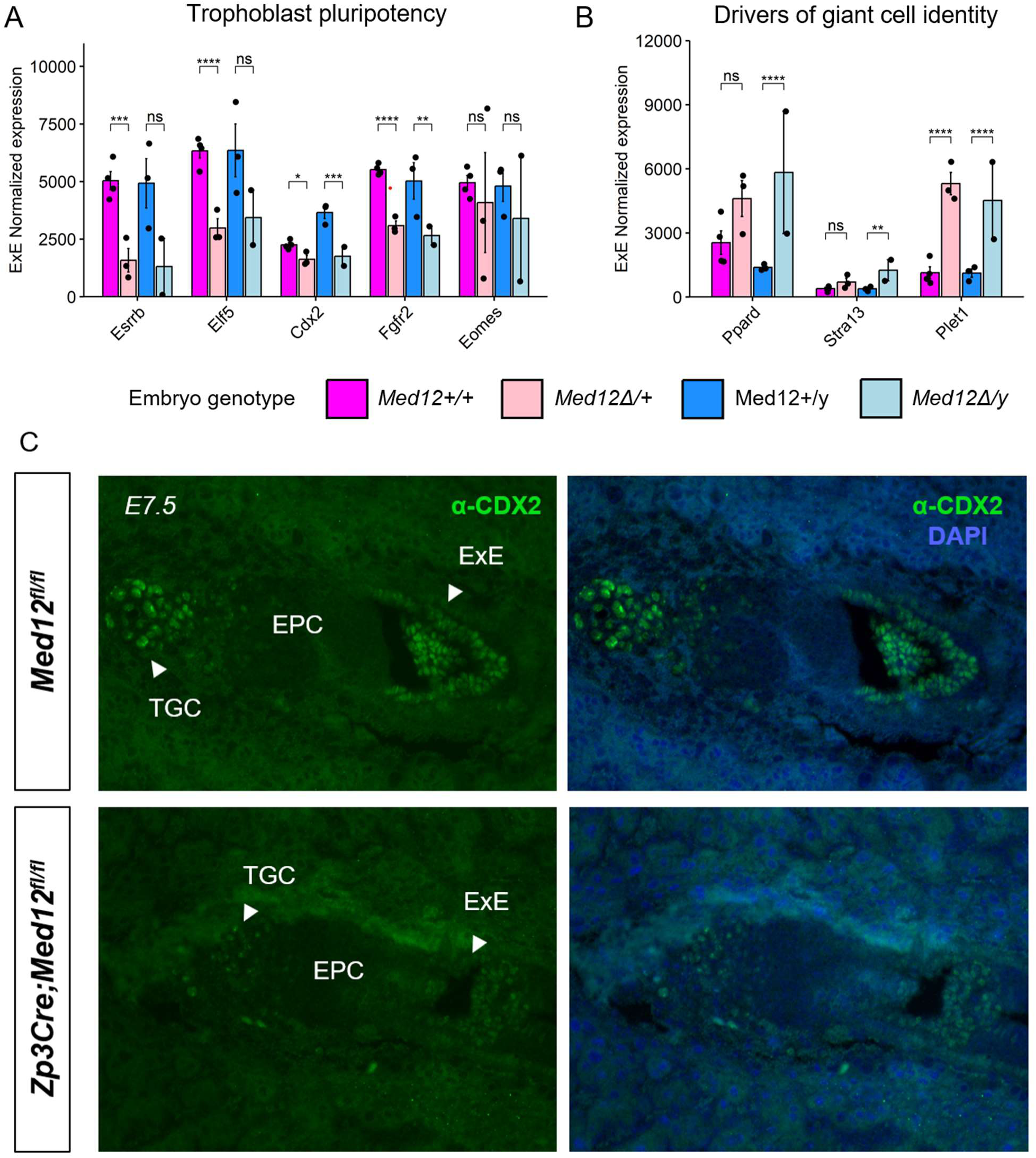
Disruption of trophoblast regulators in E7.5 embryos lacking maternal *Med12*. (A,B) Normalized expression ± S.E.M. of trophoblast pluripotency markers (A) and giant cell markers (B) in E7.5 ExE. Adjusted *p*-values from differential expression analyses (ns p≥0.05; * *p*<0.05; ** *p*<0.01; *** *p*<0.001; **** *p*<1e-4). (C) Immunofluorescence staining of E7.5 implants showing CDX2 protein (green) localized to the extraembryonic ectoderm (ExE) and primary trophoblast giant cells (TGC). Nuclei stained blue with DAPI. Implants collected from control (*Med12^fl/fl^*) and mutant (*Zp3Cre;Med12^fl/fl^*) dams that were mated with wild-type males. Representative images at 10x taken with an epifluorescence microscope.

**Table 1.**
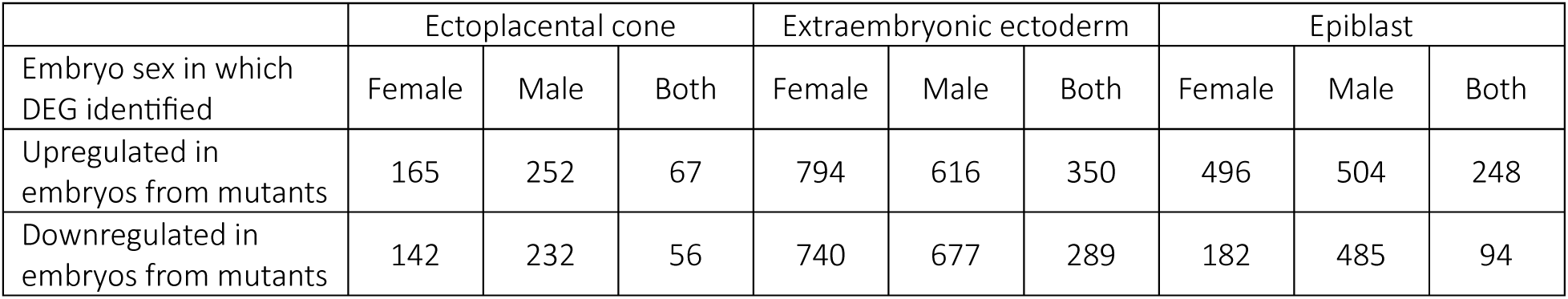
Differentially expressed genes (DEG) between E7.5 embryos from control (*Med12^fl/fl^*) and mutant (*Zp3Cre;Med12^fl/fl^*) dams (adj *p* < 0.01). Pairwise comparisons conducted for each embryonic tissue and sex. DEG grouped based on whether they were identified only when comparing males, only females, or in both sexes.

### Embryonic expression of a Med12 transgene compensates for loss of maternal Med12

Our previous research showed that *Zp3Cre;Med12^fl/fl^* females were sterile when mated with wild-type stud males (X. Wang et al., 2017). However, the *Med12* locus is X-linked, and this mating strategy did not account for the effects of paternal X chromosome inactivation (XCI), a phenomenon unique to rodents. In female mouse embryos, the paternal X chromosome is exclusively inactivated until the blastocyst stage, at which point paternal XCI is maintained in the trophectoderm but reversed in the inner cell mass (ICM), such that random XCI occurs in the embryo proper (Furlan & Galupa, 2022). Therefore, matings between *Zp3Cre;Med12^fl/fl^* dams and wild-type males would result in female embryos with a null *Med12* allele from the dam and an inactivated *Med12* allele from the sire, essentially producing a maternal-zygotic *Med12* double knock-out in all tissues except those derived from the ICM. This scenario is consistent with the low expression of *Med12* in both male (*Med12Δ/y*) and female (*Med12Δ/+*) E7.5 embryos from mutant dams, as well as the slightly higher expression in mutant female epiblast, which is derived from the ICM (Supplementary Figure 3B). In this case, the role of oocyte-derived *Med12* cannot be specifically investigated without simultaneously compromising embryonic *Med12* expression.

To address the complication of paternal XCI, we generated transgenic mice that express *Med12* from an autosome, as we had done previously with a mutant version of *Med12* commonly observed in uterine tumors (Mittal et al., 2015). Briefly, the *Med12* construct was knocked into the *Rosa26* safe harbor locus on chromosome 6. The *Med12* transgene (*ROSA^Med12^*), driven by a T7 promoter, was designed such that it could be transcriptionally activated (*ROSA^Med12(A)^*) through Cre-mediated recombination (Figure 6A). Using this strategy, we obtained stud males that express *Med12* from an autosome (Figure 6B) and mated them with mutant (*Zp3Cre;Med12^fl/fl^*) and control (*Med12^fl/fl^*) dams to test whether embryonic expression of the autosomal *Med12* transgene could compensate for the loss of maternal *Med12* (Figure 6C). As previously observed, mutant dams were infertile when mated with wild-type males; however, when mated with males that express the *Med12* transgene (ROSA^Med12(A)/+^), mutant dams produced litters at a similar frequency to control dams (Figure 6D), although litter size was significantly smaller (Figure 6E). Every pup born to a (*Zp3Cre;Med12fl/fl*) dam and (*ROSA^Med12(A)/+^*) sire carried a *ROSA^Med12(A)^* allele and an endogenous *Med12* deletion (n=29/29 pups), whereas only half of pups born to (*Med12^fl/fl^*) dams inherited the *ROSA^Med12(A)^* allele (n=16/29 pups), and all carried the intact *Med12^flox^* allele (Figure 6F). This observation explains why the average litter size for mutant dams (3.67 pups) was about half of that observed in control dams (6.17 pups), as 50% of embryos in mutant dams would not inherit the *ROSA^Med12(A)^* allele and would presumably perish around E12.5. Surprisingly, pups born to (*Zp3Cre;Med12^fl/fl^*) dams and (*ROSA^Med12(A)/+^*) sires were both viable and fertile (Supplementary Figure 6A), with no apparent phenotypic consequences resulting from *Med12* transgene expression (Supplementary Figure 6B). Moreover, *Med12* expression in (*ROSA^Med12(A)/+^*) stud males and their male offspring (*Med12Δ/y;ROSA^Med12(A)/+^*) was significantly higher than in wild-type counterparts (Supplementary Figure 6C), suggesting that mice are highly robust to *Med12* overexpression and/or that post-transcriptional mechanisms exist to control MED12 protein levels. Overall, embryonic expression of the *Med12* transgene was sufficient and necessary to rescue development of embryos lacking endogenous maternal *Med12*, indicating that the sterility of *Zp3Cre;Med12^fl/fl^* dams likely results from the persistent inactivation of the paternal *Med12* allele, rather than loss of oocyte-derived *Med12*.

**Figure 6.**
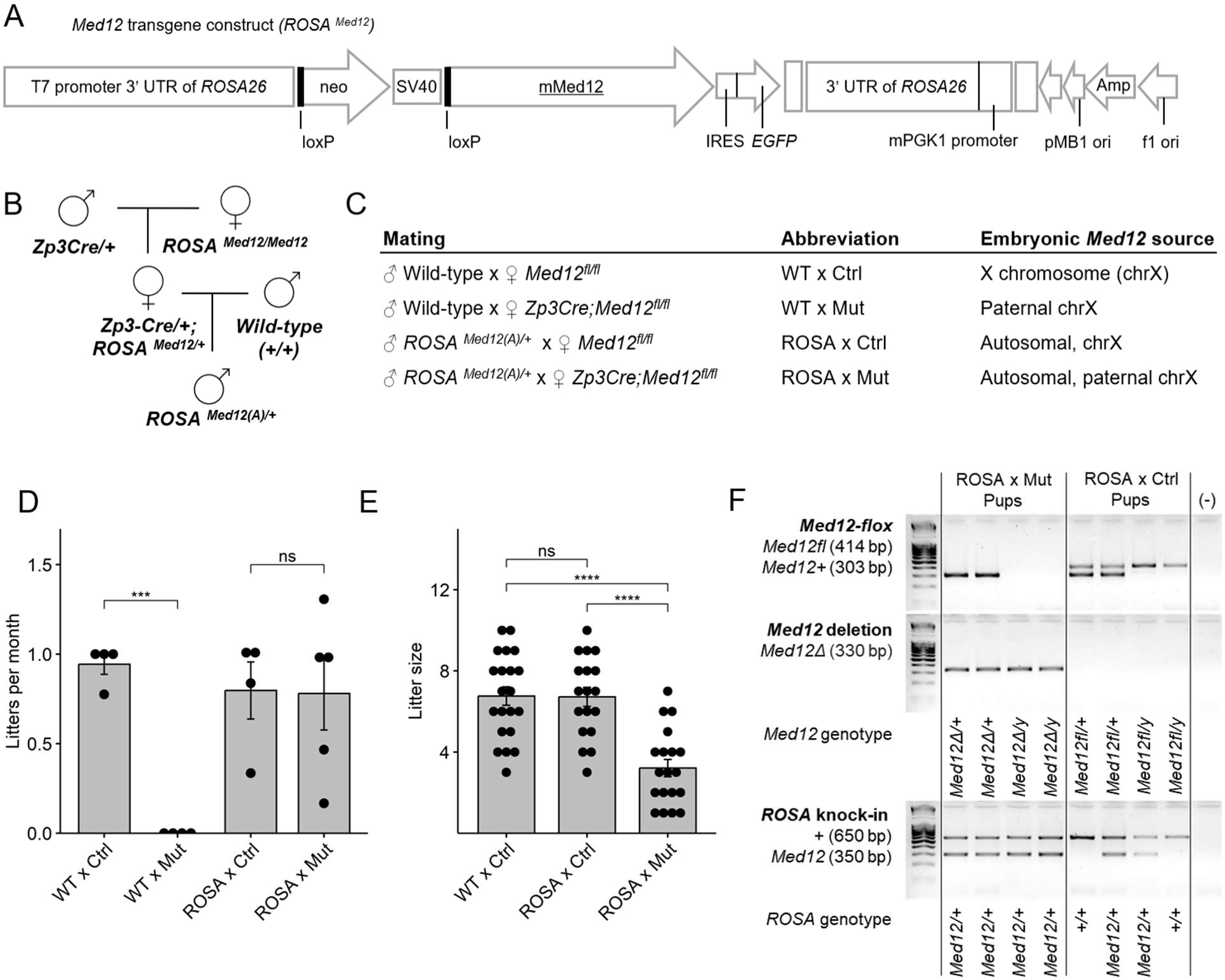
Embryonic expression of a *Med12* transgene compensates for loss of maternal *Med12*. (A) *Med12* transgene construct for knock-in to the *Rosa26* safe harbor locus on chromosome 6. (B) Breeding strategy to obtain stud males that express the activated *Med12* transgene (e.g., *ROSA^Med12(A)^*). (C) Mating strategies to test fertility of control (*Med12^fl/fl^*) and mutant (*Zp3Cre;Med12^fl/fl^*) dams, and corresponding source(s) of *Med12* in resultant embryos. (D) Litters produced per month ± S.E.M. for each breeding pair and (E) litter size ± S.E.M. for each mating combination. One-tailed unpaired Student’s t-test, adjusted by Holm-Bonferroni method (ns p≥0.05; * *p*<0.05; ** *p*<0.01; *** *p*<0.001; **** *p*<1e-4). (F) Representative genotyping of pups produced from matings between males with an active *Med12* transgene and control or mutant dams.

**Supplementary Figure 6.**
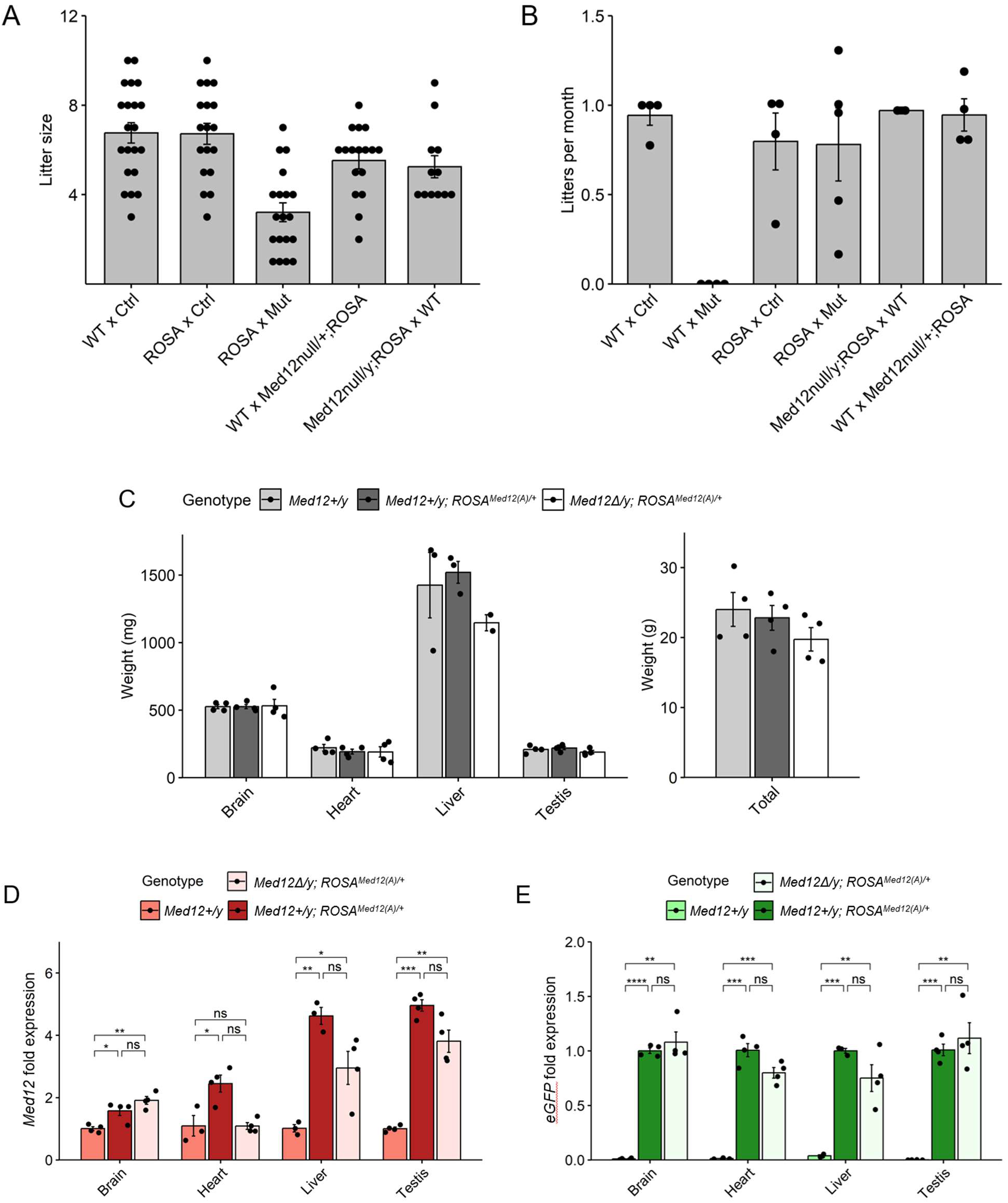
Viability of animals carrying an activated *Med12* transgene. (A) Litter size and (B) average litters produced per month ± S.E.M. for each tested mating combination (male x female). WT: wild-type, Ctrl: *Med12^fl/fl^*, Mut: *Zp3Cre;Med12^fl/fl^*, ROSA: *ROSA^Med12(A)/+^*. (C) Organ and total carcass weight ± S.E.M. of 10-week-old males that express endogenous *Med12* (*Med12+/y*), both endogenous and transgene *Med12* (*Med12+/y;ROSA^Med12(A)/+^*), or only transgene *Med12* (*Med12Δ/y;ROSA^Med12(A)/+^).* No significant differences in weight between genotypes were observed (*p*≥0.05). Two-tailed unpaired Student’s t-test, adjusted by Holm-Bonferroni method. (D,E) Relative abundance ± S.E.M. of (D) *Med12* and (E) *eGFP*, produced as a byproduct of *Med12* transgene expression, in brain, heart, liver and testis. Calculated by the ΔΔCt method using the reference gene *Gapdh*. Two-tailed unpaired Student’s t-test, adjusted by Holm-Bonferroni method (ns *p*≥0.05; * *p*<0.05; ** *p*<0.01; *** *p*<0.001; **** *p*<1e-4).

## Discussion

The study of maternal effect genes holds immense significance in the field of reproductive medicine, as disruption of these genes is catastrophic for embryonic development. A number of maternal effect genes are known to disrupt human embryonic development and are a known cause of infertility (Mitchell, 2022). Nonetheless, relatively few MEG have been identified in mammals. Previous research suggested that *Med12* functions as an MEG in mice (X. Wang et al., 2017), but the role of *Med12* in oocytes and embryos remained unexplored. Herein, we show that *Med12* is not involved in transcription regulation in the oocyte but is required in embryos for maintenance of trophoblast pluripotency and balanced differentiation of placental cell types. At E7.5, loss of *Med12* expression resulted in downregulation of pluripotency markers (*Esrrb*, *Cdx2*, *Elf5*, *Eomes*) in the ExE (Donnison et al., 2005; Gao et al., 2019; Strumpf et al., 2005). In trophoblast stem cells (TSC), which are considered analogous to the ExE (Adachi et al., 2013; Tanaka et al., 1998), the knockdown of any one of these factors triggers differentiation and upregulation of TGC markers (Latos, Goncalves, et al., 2015; Latos, Sienerth, et al., 2015). *Esrrb*-null embryos die of placental malformations, including overabundance of TGC (Luo et al., 1997), a phenotype similar to that observed in E9.5 embryos depleted for maternal *Med12*. Concurrent to the downregulation of pluripotency markers in ExE*, Med12* depletion also led to upregulation of TGC markers (*Stra13*, *Ppard*, *Plet1*). Ectopic expression of any of these factors in TSC induces terminal differentiation into giant cells (Barak et al., 2002; Hughes et al., 2004; Murray et al., 2016; Nadra et al., 2006). Notably, higher *Plet1* expression in TSC accelerates differentiation towards TGC, whereas low *Plet1* favors syncytiotrophoblast fate (Murray et al., 2016), potentially explaining both the overabundance of TGC and the diminished syncytiotrophoblast layer observed in maternal *Med12*-depleted placenta. Altogether, we conclude that loss of *Med12* disrupts pluripotency maintenance in the ExE and skews differentiation towards the TGC fate.

This previously undescribed role of *Med12* in trophoblast pluripotency maintenance is consistent with reports that MED12 co-localizes with histone acetylases at super enhancers in TSC (Lee et al., 2019). Moreover, *Med12* knock-down (KD) in other types of stem cells similarly disrupts pluripotency maintenance. In embryonic stem cells (ESC), Mediator and cohesin coordinate 3D interactions between ESC-specific enhancers and pluripotency genes; *Med12*-KD abolishes these interactions, leading to downregulation of pluripotency markers (Kagey et al., 2010; Phillips-Cremins et al., 2013; Whyte et al., 2013). Similarly, in hematopoietic stem cells (HSC) MED12 stabilizes histone acetylases at HSC-specific enhancers; *Med12*-KD leads to enhancer deacetylation and loss of stemness *in vitro*, and bone marrow HSC depletion and lethality *in vivo* (Aranda-Orgilles et al., 2016). Together, these studies suggest that *Med12* plays a key role in pluripotency maintenance across many different stem cell lineages. Future studies will elucidate the mechanisms that govern cell type-specific *Med12* activity (e.g., binding to TSC-versus ESC-specific super enhancers), its relation to the epigenome and chromatin conformation, and whether *Med12* acts independently of the core Mediator complex in the context of pluripotency.

In our efforts to specifically interrogate the function of maternal *Med12*, we found that embryos from *Zp3Cre;Med12^fl/fl^* dams had persistently low levels of *Med12*, even in the presence of a functional paternal copy. We hypothesized that programmed paternal XCI compromises embryonic *Med12* expression, effectively generating a maternal-zygotic double knock-out. Using a knock-in approach, we showed that expression of an autosomal *Med12* transgene rescues embryonic development from *Med12*-depleted oocytes. This finding indicates that the absence of *Med12* in the oocyte does not engender any irreversible effects, either transcriptomic or epigenetic (e.g., imprinting), that cannot be corrected through embryonic *Med12* expression. This study highlights the difficulties involved in studying the developmental roles of X-linked genes in the mouse, which likely explains why there are no X-linked MEG described to date in the mouse. Although *Med12* may not fit the strict definition for MEG in mice, its role in oogenesis and development in non-rodent mammalian species remains to be investigated. Overall, we conclude that oocyte-derived *Med12* is required for murine development because of the rodent-specific phenomenon of programmed paternal X chromosome inactivation, and that *Med12* is essential for pluripotency maintenance in the trophoblast, as well as proper differentiation and morphology of the placenta and embryo proper.

## Methods

### Experimental mice

All procedures were approved by the IACUC of the University of California, San Francisco (UCSF) and conducted in accordance with NIH and UCSF guidelines for the care and use of laboratory animals. Mice were housed under standard conditions with a 12-hour light-dark cycle and provided ad libitum access to food and water. *Med12^flox^* mice were a gift from Dr. Heinrich Schrewe (Max-Planck Institute, Berlin, Germany) (Rocha et al., 2010), and were maintained on a C57BL/6J/129SV background. *Zp3Cre* mice were purchased from Jackson Laboratories (Stock #003651) and were maintained on a C57BL/6J background (De Vries et al., 2000). *Med12* autosomal knock-in mice were generated as previously described by subcloning full-length *Med12* cDNA into the pROSA26-DV1 vector and inserting into the autosomal *Rosa26* locus (Mittal et al., 2015). *Med12* knock-in mice were maintained on an FVB/C57BL/6/129SV background. To test dam fertility, single females of 6 to 10-weeks of age were continuously housed with stud males of proven fertility for up to 6 months, during which time the number of litters, litter size, and sex ratios of pups were recorded. Genotyping was performed following standard PCR protocols and using DNA extracted from either tail or ear biopsies (Supplementary Table 1).

**Supplementary Table 1.**
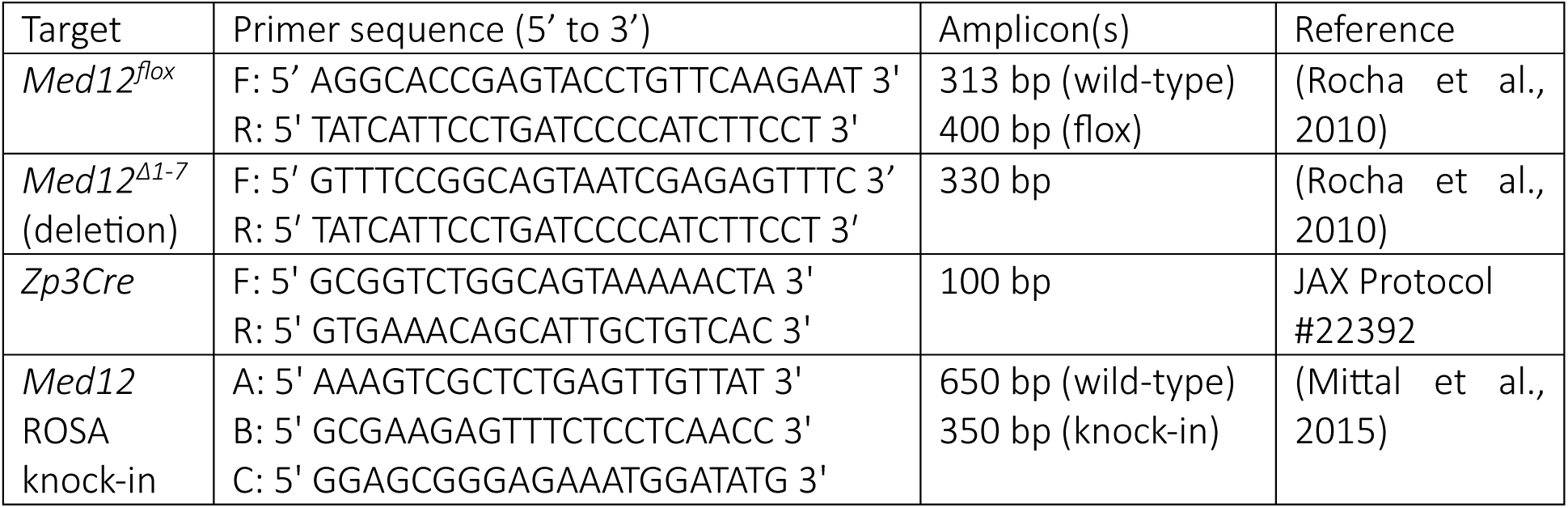
Primers used for PCR genotyping of mice.

### Histological analysis and immunofluorescence staining of uteri

Uteri harvested at various stages of gestation were fixed in 10% formalin overnight, embedded in paraffin, and serially sectioned at 5 µm thickness. Sections were stained with hematoxylin and eosin for histological analysis. For immunofluorescence staining, paraffin sections were dewaxed with xylene, rehydrated, and washed. For antigen retrieval, slides were then incubated in Tris-EDTA buffer (10 mM Tris, 1 mM EDTA, 0.05% Tween, pH 9.0) for 25 min in a steamer, incubated at room temperature for 30 min, washed twice with distilled water for 5 min, then once in 1X PBS-T for 5 min. Slides were blocked in 10% goat serum in 1X PBS-T for 1 h and then incubated with primary antibody rabbit anti-mouse CDX2 (Abcam, ab76541) at a 1/1000 dilution in 1X PBS-T overnight at 4°C. Slides were washed 3x 5 min in 1X PBS-T, then incubated with Alexa Fluor 488 goat anti-rabbit IgG (Jackson ImmunoResearch Laboratories, 115-545-003) at a 1/250 dilution in 1X PBS-T for 1 hour at room temperature. Slides were washed 3x 5 min in 1X PBS-T and mounted with ProLong Gold Antifade with DAPI (ThermoFisher, P36931). Images were captured with an epi-fluorescent microscope at 10X and processed with ImageJ (v1.50d).

### Oocyte collection for RNA-seq

Ovaries from 12 day old females were incubated in digestion media (1x DPBS, 2 mg/mL collagenase type 1, 2 mg/mL BSA) at 37°C for 1 hour, then supplemented with 40 mM EDTA (pH 8.0) and incubated at 37°C for 10 min. Digested ovaries were centrifuged at 1000 rpm for 5 min, and cell pellets resuspended in pick-up media (1x MEM-Alpha, 25 mM NaHCO_3_, 2 mg/mL BSA). Oocytes were transferred to wash buffer (1x MEM-Alpha, 25 mM NaHCO_3_, 1 mg/mL BSA), then to 1x DPBS, and finally stored in media solution at-80°C. For GV and MII oocyte collection, female mice older than 6 weeks were synchronized by 5 IU PMSG. For GV oocytes, ovaries were collected 48 h after PMSG administration and transferred to collection media (1X Hepes-MEM, 6 mM NaHCO3, 0.23 mM sodium pyruvate, 3 mg/mL BSA) with 1/1000 cilostamide. Follicles were mechanically disrupted using two needles and oocytes were selected based on size and morphology. Oocytes were washed in maturation media (1X MEM-Alpha, 6 mM NaHCO_3_, 3 mg/mL BSA) with 1/1000 cilostamide, then in 1X DPBS, and stored in Trizol solution at -80°C. For MII oocytes, female mice were injected with 5 IU hCG 48 h after PMSG, and oocytes were collected 12-15 h after hCG. Ovaries were transferred to collection media. After follicle disruption, cumulus complexes were transferred to maturation media and denuded with 0.3 mg/mL hyaluronidase for 1 min. Denuded oocytes were washed in maturation media, then 1X DBPS, and stored in Trizol solution at -80°C.

### E7.5 embryo collection for RNA-seq

Female mice older than 6 weeks were synchronized by 5 IU PMSG for 48 h, then injected with 5 IU hCG and mated with wild-type stud males of proven fertility. Uteri were collected from euthanized females 8 days post conception and transferred to HEPES-buffered DMEM supplemented with 10% FBS. Each embryo was carefully separated from the maternal decidua and dissected into the ectoplacental cone, extraembryonic ectoderm, and epiblast. Tissues were stored in Trizol solution at-80°C prior to RNA extraction.

### RNA-seq library preparation and sequencing

Total RNA was extracted from samples stored in Trizol solution by phenol-chloroform extraction. The aqueous phase was subjected to column purification using the PicoPure RNA Isolation kit (ThermoFisher, KIT0204) and treated with DNase I (Qiagen, 79254) to remove genomic DNA. For samples not stored in Trizol, total RNA was extracted using the Extraction Buffer provided in the PicoPure RNA Isolation kit. RNA integrity (RQN > 7) was evaluated with the 5200 Fragment Analyzer System (Agilent Technologies). For oocyte samples, libraries were constructed with the SMART-Seq v4 Ultra Low Input RNA kit (Takara Bio) and sequenced on the HiSeq platform to generate 50 bp single-end reads. For E7.5 embryo samples, libraries were constructed with the Tecan Universal Plus Total RNA library preparation kit (Tecan Life Sciences) and sequenced on the NovaSeqX platform to generate 50 bp single end reads.

### RNA-seq data analysis

Raw reads were trimmed with Trim_Galore (v0.6.7) and aligned to the GRCm39 assembly with STAR (v2.7.10b). Low quality alignments (q < 5) were removed with SAMtools (v1.16) and raw counts for genes in the Ensembl (v108) annotation were calculated with HTSeq (v2.0.2) in mode “intersection-nonempty”. Raw counts were log transformed with the R package DESeq2 (v1.36.0). Principal components analyses were conducted with the plotPCA function, considering the top 5,000 genes with highest variable expression. Pearson correlations were calculated with cor function from the R package stats (v4.3.3) and plotted with pheatmap (v1.0.12). Differentially expressed genes were identified with DESeq2 using an adjusted *p* value threshold of 0.01. For oocytes, pair-wise comparisons were conducted to compare samples from mutant and control dams for each stage (D12, GV, MII). For E7.5 embryos, separate pair-wise comparisons were conducted to compare samples from mutant and control dams for each embryonic tissue (EPC, ExE, EPI) and embryo sex (male or female). Volcano plots were generated with EnhancedVolcano (v1.20.0). Functional enrichment of gene sets was calculated using the DAVID webtool (v2024q1). Bar plots and bubble plots were generated with ggplot2 (v3.5.1) and ggpubr (v0.6.0).

### Oocyte collection for RT-qPCR

Female mice older than 6 weeks were synchronized by 5 IU PMSG. For GV oocytes, ovaries were collected 48 h after PMSG administration and transferred to warmed M2 media. Follicles were mechanically disrupted using two needles and oocytes were selected based on size and morphology. For MII oocytes, mice were injected with 5 IU hCG 48 h after PMSG, and oocytes were collected 12-15 h after hCG. Cumulus oocyte complexes (COCs) were isolated from the ampulla in warmed M2 media. COCs were denuded with 0.3 mg/mL hyaluronidase for 1 min. After isolation, oocytes were washed 3x in M2 media, frozen in minimal media (<5 µL) on dry ice, then stored at -80°C. Oocytes were lysed using the SingleShot Cell Lysis kit (BioRad, 172-5080). To each sample, 10 uL of lysis mix (9.6 ul SingleShot Cell Lysis buffer, 0.2 ul proteinase K, 0.2 ul DNase) was added, and samples were incubated 15 min at 22°C, 5 min at 37°C, then 5 min at 75°C. Lysates were immediately subjected to reverse transcription with the SuperScript III First-Strand Synthesis System using random primers (ThermoFisher, 18080-051).

### Adult male tissue collection

Tissues from adult male mice were harvested at 10 weeks of age. Brain, heart, liver, and testis were weighed and stored in Trizol solution at -80°C. Total RNA was extracted by phenol-chloroform and purified with the RNeasy Mini kit (Qiagen, 74104) and DNase I (Qiagen, 79254). RNA extraction from brain was modified as previously described (Dzhala et al., 2022). Briefly, after chloroform extraction an equal volume of isopropylalcohol was added to the aqueous phase, and samples were centrifuged to pellet total RNA, which was resuspended in 70% ethanol before purification with the Qiagen RNeasy Mini kit. Purified RNA was subjected to reverse transcription with the SuperScript III First-Strand Synthesis System using random primers (ThermoFisher, 18080-051).

### Real-time quantitative PCR (RT-qPCR)

Quantitative PCR was performed on the StepOnePlus instrument (ThermoFisher) using the SsoAdvanced Universal SYBR Green Supermix (Biorad, 1725271) (Supplementary Table 2). C_T_ values were normalized to *Gapdh* and relative transcript abundance was determined with the ΔΔC_T_ method.

### Statistics

Student’s *t* tests were conducted with the t_test function from the rstatix package (v0.7.2). *P* values were adjusted for multiple testing using the Holm-Bonferroni method. An adjusted *P* value less than 0.05 was considered statistically significant.

**Supplementary Table 2.**
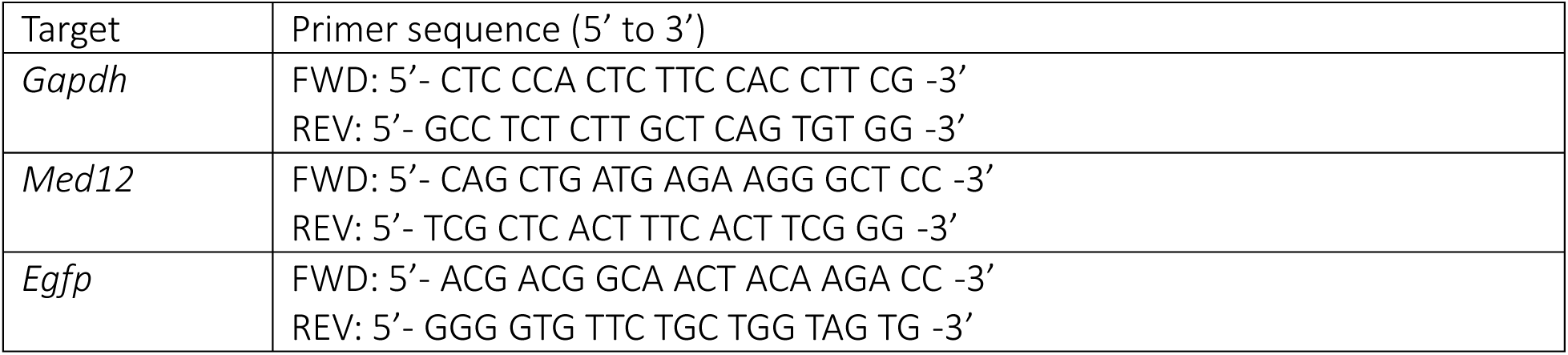
Primers for RT-qPCR.

## Acknowledgments

The authors greatly appreciate Dr. Heinrich Schrewe for donating the mice. The authors thank the University of California San Fransisco Genomics CoLab for assistance in library preparation and sequencing. MMH was supported by NIH grant 5T32HD7263-39.

## Notes

### Competing Interest Statement

The authors have declared no competing interest.

